# Overexpression of *EiKCS* confers paraquat-tolerance in rice *(Oryza sativa* L.) by promoting polyamine pathway

**DOI:** 10.1101/2020.12.11.421701

**Authors:** Qiyu Luo, Shu Chen, Jiazheng Zhu, Laihua Ye, Nathan Daniel Hall, Suma Basak, J. Scott McElroy, Yong chen

## Abstract

Paraquat is an important bipyridine herbicide by acting on the photosynthetic system of the plants and generating reactive oxygen species leading to cell death, whereas the mechanism of the paraquat resistance remains to be explored. In this study, a putative paraquat-resistant gene *EiKCS* from goosegrass *(Eleusine indica* L.) was isolated and overexpressed in a transgenic rice *(Oryza sativa* L.). This transgenic rice (KCSox) was treated by exogenous spermidine and paraquat and then was analyzed by qualitative and quantitative proteomics. Overexpressing of *EiKCS* enhanced paraquat tolerance in KCSox by the accumulation of endogenous polyamines whose dominant presences of polyamines benzoylation derivatizations in rice were C_18_H_20_N_2_O_2_, C_28_H_31_N_3_O_3_, and C_38_H_42_N_4_O_4_. The mechanism underlying the improving tolerance enhanced antioxidant capacity of ROS systems and light-harvesting in photosynthesis in KCSox rice leaves to reducing paraquat toxicity. The protein β-Ketoacyl-CoA Synthase (EiKCS) encoded by the *EiKCS* gene promoted the synthesis and metabolism of proteins of the polyamine pathway. Three cofactors CERs were identified and positively correlated with the function of EiKCS on very-long-chain fatty acids (VLCFAs) biosynthesis via promoting the polyamine pathway and inhibiting the links with the TCA pathway and fatty acid pathway to responding to the paraquat tolerance in the KCSox rice, which also caused the prolongation of the overproduction of spermine and a transient increase of intracellular malondialdehyde (MDA). These results expanded the polyamines pathway manipulated in cereals using genetic engineering to clarify the mechanism of paraquat-tolerance.

**One Sentence Summary:** A putative paraquat-resistant *EiKCS* gene from the goosegrass overexpressing in the rice resulted in the accumulation of polyamines, especially the spermine, and promoted the proteins in polyamine pathways by its EiKCS protein under paraquat stress.

## Introduction

Paraquat (1,1’-dimethyl-4,4’-bipyridinium dichloride) as one of photosystem I (PSI) inhibitors are worldwide used in agriculture and gardening because it rapidly kills weeds and green plants (Li et al., 2013). In green plants, paraquat inhibits normal functioning of PSI by stopping electron transport to cause cell membrane disruption and plant death for paraquat ion PQ^2-^ accepting electron to form paraquat monocation free radical PQ^2-^ (Gao et al., 2018). In this manner, it can result in the production of toxic reactive oxygen species (ROS) including H_2_O_2_, HClO and free radicals O^2-^ to facilitates the killing effect of paraquat (Dong et al., 2016). In addition to its weed control efficacy, paraquat is unsubstituted in bipyridine herbicides for its timely inactivation upon reaching the soil1(Reczek et al., 2017).

During single repeatability applications of commercial paraquat for decades, there are more than 30 species of paraquat-resistant weeds have been reported worldwide (http://www.weedscience.org). And the mechanism of resistance to paraquat was recognized as a dominant (or semi-dominant) single gene trait and was also involved in vacuolar sequestration (Qin et al., 2010, Hawkes 2014, Caio et al., 2017). Goosegrass *(Eleusine indica.* L) as one of the most serious invasive weeds in crops on many countries due to its paraquat-resistance (Chuah et al., 2010; Jalaludin et al., 2015; Chen et al., 2017). But the resistance and detoxification mechanism of in paraquat-resistant biotypes of goosegrass is considered to be related to polyamines (An et al., 2014; Deng et al., 2019; Luo et al., 2019).

Polyamines are proposed to be specific structural similarities with paraquat in **Figure 1** (Dinis-Oliveira et al., 2008; Palmer et al., 2010; Blanco et al., 2014; Fujita et al., 2014). Polyamines are essential for survival and growth of all living organisms, including putrescine (Put), spermidine (Spd), and spermine (Spm) as the major polycationic compounds in plants. And polyamines are not only involved in cell death, seed germination and root formation but also ROS stress pathways induced by paraquat (Mulangi et al., 2012a; Mulangi et al., 2012b; Tanou et al., 2014). The polyamine biosynthetic pathway in plant is clearly clarified that arginine synthesizes putrescine via ornithine or N-carbamoylputrescine and putrescine successively converts to spermidine and then spermidine to spermine (Alcázar et al., 2010; Naka et al., 2010; Takahashi et al., 2018). Despite this progress, it remains unknown how paraquat regulated the polyamine pathway and cross-talk with other stress pathways.

**Figure 1.**
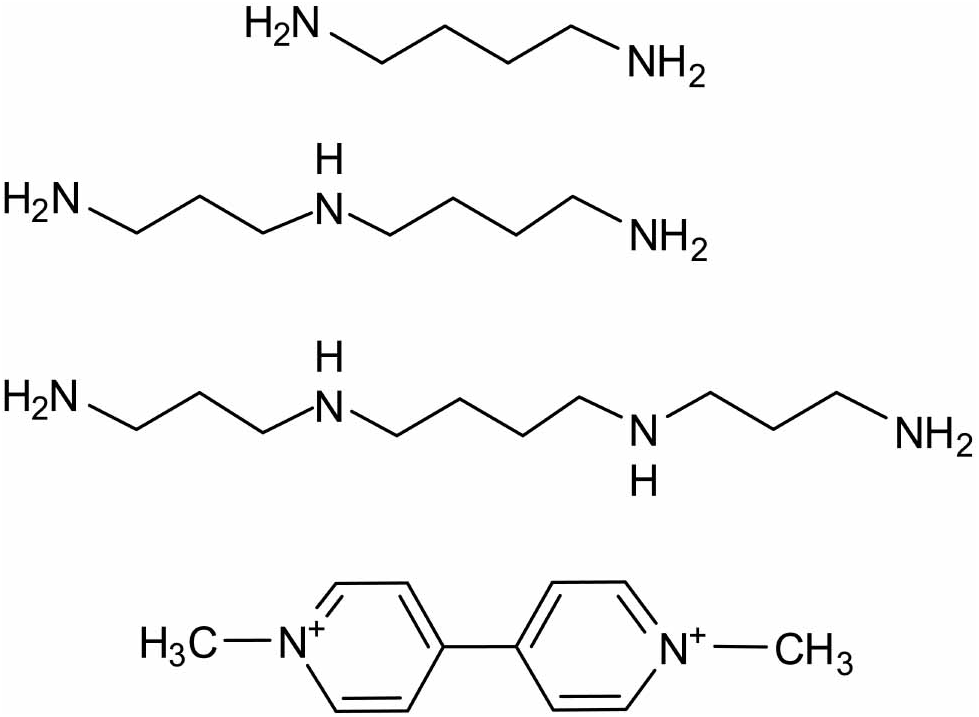
The polyamines and paraquat: (from top to bottom) putrescine, spermidine, spermine and paraquat (1,1’-dimethyl-4,4’-bipyridinium dichloride).

In this study, a putitave paraquat-resistant gene *β-ketoacyl-CoA synthase* (*EiKCS*) is cloned from paraquat-resistant goosegrass and overexpressed in rice (Oryza sativa) to investigate new resistant-site of action about non-selective activity in genetically modified crops (GMO). The results showed that paraquat-tolerance in KCSox rice was enhanced by promoting the biosynthesis of endogenous polyamines and encoding protein EiKCS to regulate proteins in polyamine degradation pathway and cross-talk with fatty acid, VLCFAs and TCA cycle, including a special eceriferum protein (CER) as cofactors. The aim of this study was to provide useful insight for future gene function studies in more non-selective herbicide resistance in crops.

## Results

### Construction and identification of the KCSox rice

A significantly increased transcript level of the KCS gene, which was previously named as the *PqE* gene, was observed in leaves of paraquat-susceptible and paraquat-resistant biotype goosegrass in 90 min after paraquat spraying treatment (Luo et al, 2019). The coding sequences (CDS) of KCS genes from the two goosegrass biotypes were aligned and no non-synonymous variation was identified between the sequences (**Supplemental Fig. S1**). The phylogenetic analysis showed that *EiKCS* was closest to KCS gene of rice (Oryza_sativa_subsp_indica_A2X281) (**Fig. 2A**). Thus, the CDS sequence of EiKCS (1551bp) was amplified from susceptible goosegrass (**Fig. 2B**) and overexpressed in a male-fertile japonica (O.sativa ssp. Japonica L.) variety rice Zhonghua11 (WT). The KCSox rice was driven by Ubiquitin promoter of pCUbi1390 vector about 10860bp (**Fig. 2C**). The total DNA was separately extracted from a large number of leaves samples in KCSox rice by a simplifying method TPS (**Supplemental Fig. S2**). A single PCR amplification target band appear in the positive identification of each generation of KCSox (**Fig. 2D**). In order to better identify and screen the positive in later generations, KCSox and WT seeds germinated in the culture dishes and soils after soaking in a hygromycin solution with the same concentration gradient (**Fig. 3A and 3B**). Both culture methods distinguished KCSox and WT at a low concentration of hygromycin but culture in dishes was better than in soil for its less effect on germination rate (**Fig. 3C and 3D**). Therefore, KCSox seeds germinating in the culture dishes after soaking in hygromycin solution (25mg/L) for 2 days could eliminate the interference of WT seeds (**Fig. 3E**), which also could apply for GMO progenies with the *Hygromycin* marker gene. In addition, exogenous hygromycin was not applicable on rice leaves to screening because it inhibited the nutritional growth of rice and caused damages on leaves (**Fig. 3F and 3G**).

**Figure 2.**
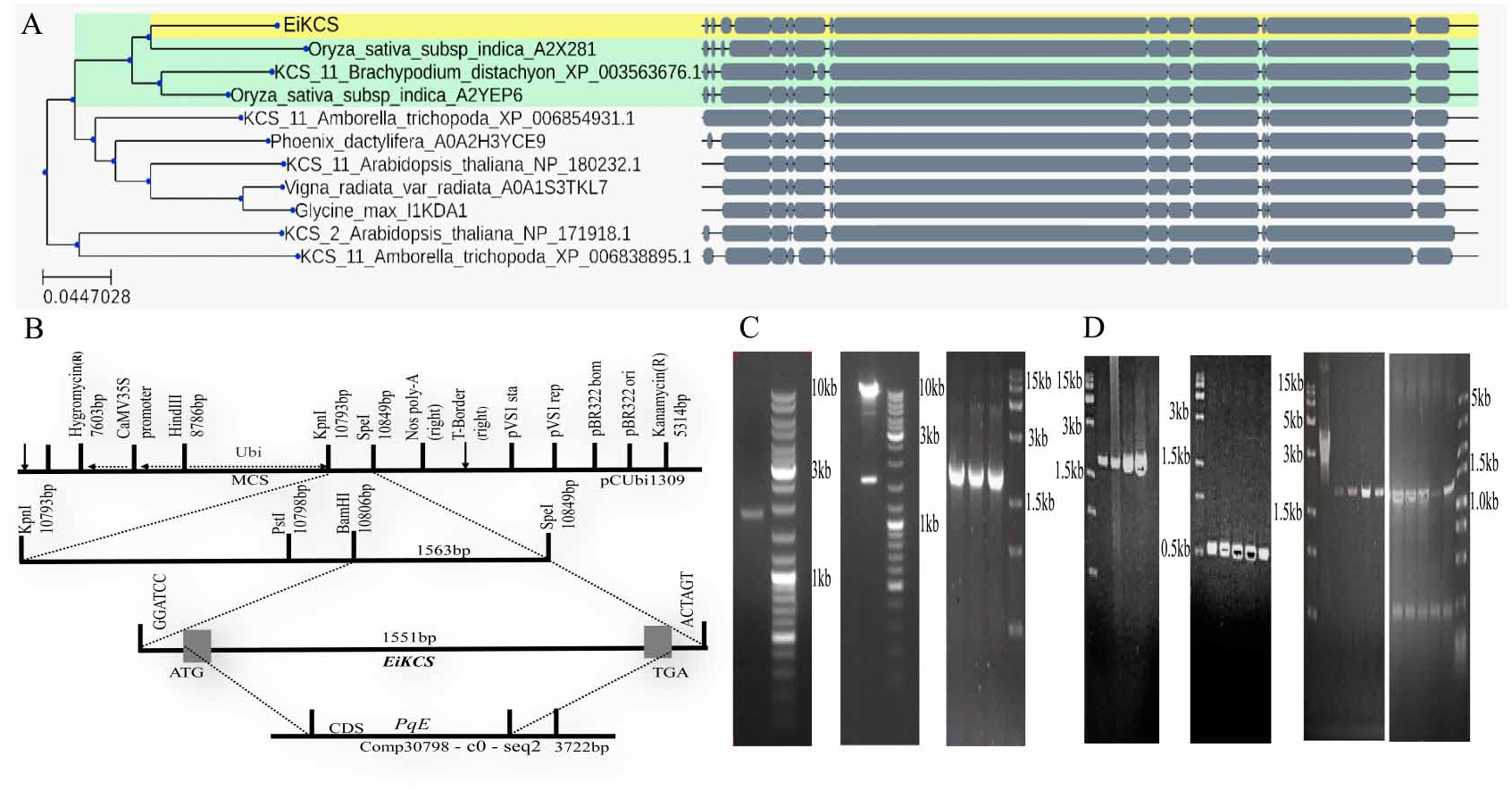
Phylogenetic analysis, construction of overexpression vector, and positive identification of transgenic T0-T3 generations rice. **A.** Transcriptomes sequence of *EiKCS* gene with all homogeneous genes in the NCBI database and its evolutionary tree. Transcriptomes sequence of *EiKCS* gene is colored yellow, and homologous gene sequence with the closest relationship is colored green. **B.** Genetic mapping of *EiKCS.* Markers used for vector plasmid structure with bases numbers are shown on the top, and the length of the gene sequence and the modification of the enzyme site are shown in the amplification section. **C.** PCR amplified the construction of overexpression vector. CDS sequence of EiKCS gene (1551bp) was amplified by primer PqE-F/R from goosegrass. The sequence of *EiKCS* gene with BamHI and Spel sites (1563bp) was double-digested from overexpression pCUbi1309::EiKCS vector. The target sequence of *EiKCS* gene(1898bp) was amplified by primer pCUbi1390-F/R from *Agrobacterium tumefaciens* Mono-clones. **D.** The target sequence of *EiKCS* gene(1898bp) was separately amplified by primer pCUbi1390-F/R from former and latter 2 samples of T0 and T1 generations transgenic rice. The target sequence of *Hygromycin* marker gene (289bp) was separately amplified by primer Hyg-F/R from from former and latter 2 samples of T0 and T1 generations KCSox. The target sequence of *EiKCS* gene(1898bp) was amplified by primer pCUbi1390-F/R from T2 generations KCSox, a template of pCUbi1309::PqST3 vector as control on first line (3413bp). The target sequence of *EiKCS* gene (1468bp) was amplified by primer PqE-1-F/R from T3 generations KCSox.

**Figure 3.**
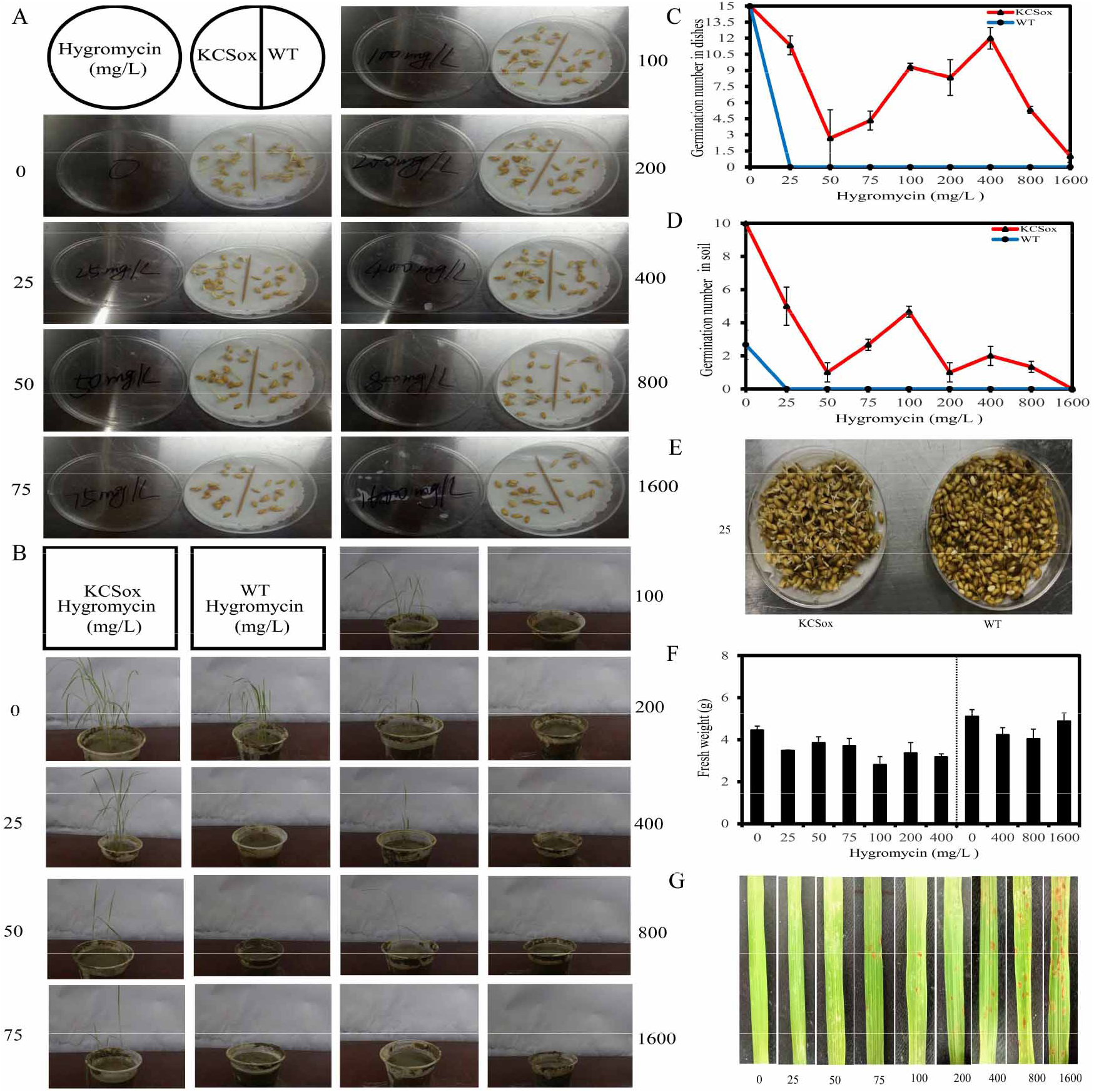
Hygromycin screened KCSox rice progenies. **A** and **B.** Germination phenotypes in culture method in dishes and soils by using ddH_2_O showed KCSox rice (oval) on the left and WT (RHY1179) a commercial RHY1179 wild-type rice (long) on the right after soaking in 0, 25, 50, 75, 100, 200, 400, 800 and 1600 mg/L for 2 days. **C** and **D.** Germination number of KCSox and WT (RHY1179) rice in the culture dishes and soils after soaking 2 days with the gradient of hygromycin solutions (3 replicates). **E**. Germination of T2 generation of KCSox and WT (Zhonghua11) seeds after soaking in 25mg/L Hygromycin solution for 2 days. KCSox and WT (Zhonghua11) were the same genetic background materials. **F.** The above-ground fresh weight of each pot was measured on the 7th day after sprayed with the gradient of hygromycin solutions (3 replicates). **G.** The leaves of WT (RHY1179) were sprayed with the gradient of hygromycin solutions. Photographs were taken as a record on 7th days.

### KCSox or pre-applied exogenous spermidine confer tolerance to paraquat

Spraying paraquat can cause visually withering and dying of WT leaves at 6-9 leaf stage from 810mg·L^−1^ (**Fig. 4A**). To measure median effective dose of paraquat (ED50), two gradients of paraquat solution at 0, 10, 30, 90, 270, 810, 2430mg/L and then 0, 90, 180, 270, 360, 450, 630, 810 mg·L^−1^. The results showed the inhibition rate reached 54.42% at 270 mg·L^−1^ (**Fig. 4B**). And next the 270 mg·L^−1^ paraquat was used instead of ED50 to apply on spraying both WT and KCSox lines at the 6-9 leaf stage. KCSox seedlings showed tolerance to paraquat while the growth of WT was significantly inhibited after spraying (**Fig. 4C**). Their fresh weight of above ground WT had a significant 32.22% decreased of as compared with control (P<0.05) while KCSox had an 19.57% decreased of fresh weight of above ground as compared with control (**Fig. 4D**). Moreover, exogenous low concentration spermidine could promote the growth of WT leaves at 6-9 leaf stage and reach its maximum biomass at 1.5 mmol/L (**Fig. 4E**). And pre-applied exogenous spermidine alleviated the damage of paraquat to WT (**Fig. 4F**). The 12 hours is the best time for pre-applying 1.5 mmol/L spermidine to alleviate the damage of 270 mg·L^−1^ paraquat with the fresh weight of above ground decreased the least (**Fig. 4G**). Therefore, these optimal treatment conditions were used to further study the mechanism of paraquat-tolerance in KCSox relating polyamines, ROS, MDA, Fv/Fm and so on.

**Figure 4.**
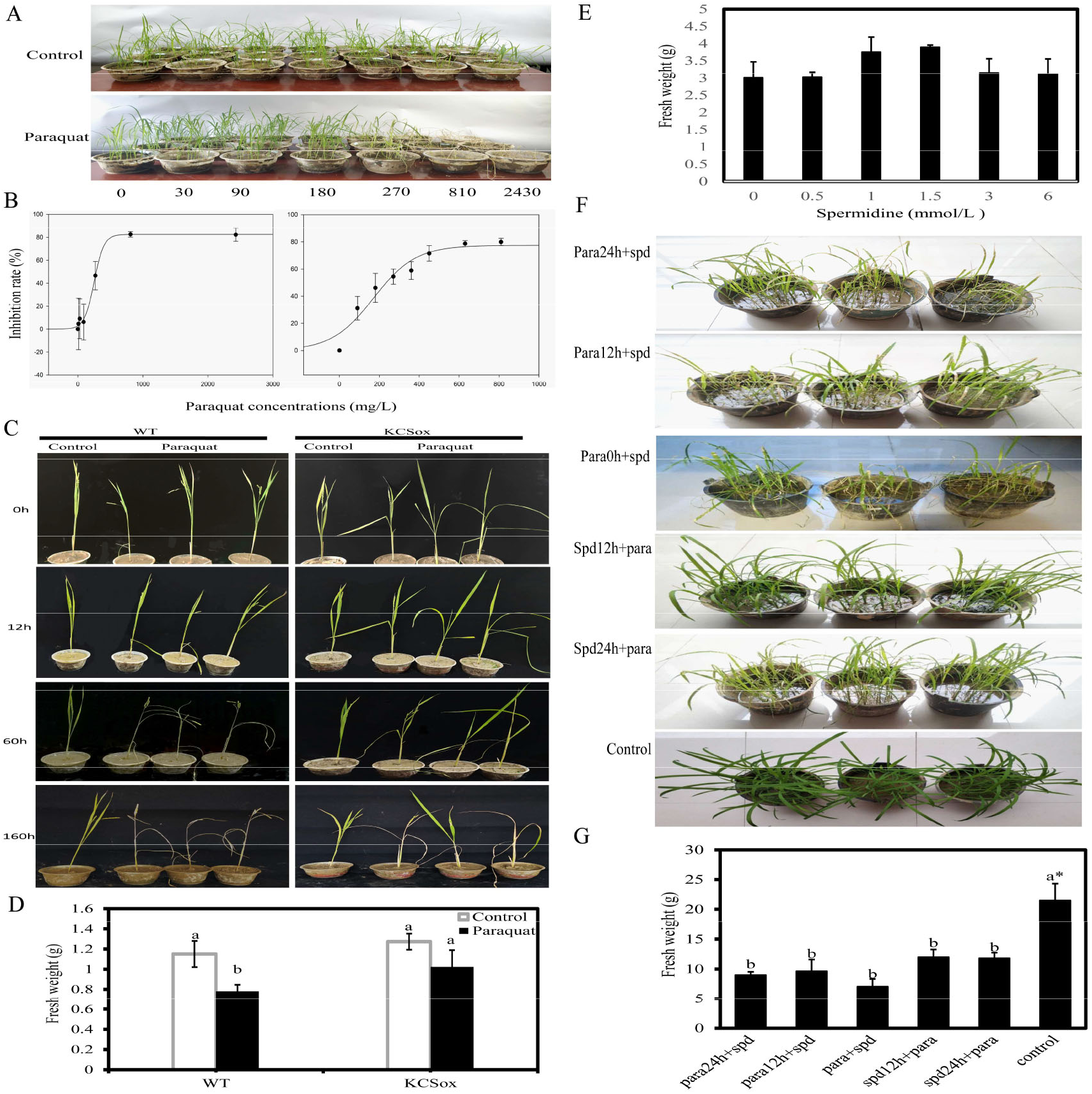
Growth indexes of KCSox and WT under paraquat and spermidine treatments. **A.** The phenotypes of paraquat damages of WT(RHY1179) leaves at 6-9 leaf stage with sprayed paraquat concentration gradient 0, 10, 30, 90, 270, 810, 2430mg/L on 4th days. **B.** Inhibition rates of WT(RHY1179) at 6-9 leaf stage were sprayed paraquat concentration gradient 0, 10, 30, 90, 270, 810, 2430mg/L, and 0, 90, 180, 270, 360, 450, 630, 810mg/L (Each 3 replicates) on 7th days. **C.** The phenotypes of paraquat damages of KCSox and WT (zhonghua11) at 6-9 leaf stage after spraying 270mg/L paraquat solutions. **D.** Fresh weight of KCSox and WT (zhonghua11) at 6-9 leaf stage after spraying 270mg/L paraquat solutions on 7th days. Each had 9 replicates under the paraquat treatment and 3 replicates under the control (ddH_2_O). Different letters indicated different significance (P < 0.05) in intra-group comparison. **E.** Fresh weight of WT (RHY1179) sprayed spermidine solutions with 0.5, 1.0, 1.5, 3.0 and 8.0 mmol/L is compared to the control (ddH_2_O) on 7th days. **F.** The phenotypes of WT (RHY1179) leaves at 6-9 leaf stage on two days after spraying 270 mg/L paraquat with 1.5mmol/L spermidine pretreatment in different time. Para24h+spd indicated spraying paraquat 24h after spermidine spraying. Para12h+spd indicated spraying paraquat 12 h after spermidine spraying. Para0h+spd indicated spraying paraquat immediately after spermidine spraying. Spd12h+para indicated spraying paraquat 12h before spermidine spraying. Spd24h+para spraying paraquat 24h before spermidine spraying. Control indicated spraying ddH_2_O. Different letters indicated different significances (P < 0.05) and an asterisk indicated significant differences (P < 0.01) among all treatments.

### Overexpression of *EiKCS* gene increases endogenous polyamines in KCSox

After benzoylation, the standards of putrescine, spermidine and spermine were distinctly separated by the used conditions with their retention time respectively (**Fig. 5A**). Total free polyamines extracted from KCSox and WT rice leaves were similarly derivatized and analyzed by HPLC. The peaks of polyamines standards and rice samples keep the consistence of the retention time at ca. 3.112 min (Put), 3.910min (Spd) and 4.414 min (Spm). To confirm this, we made a progress on the quantitative analysis of rice polyamines by HPLC-MS/MS. The results clearly showed the exact mass of polyamine derivatization products were 297.2 (Put), 458.2 (Spd) and 619.3 (Spm) in both standards and rice samples (**Fig. 5B**). To further validate our method, all the possible molecular formulas of polyamines benzoylation derivatizations are analyzed. Due to the amidation reaction is a classical reaction, each N site of polyamines may bind benzene ring when acyl chloride reacts with amide completely. Thus, there are seven possible molecular structures for polyamines derivatizations with two for putrescine, three for spermidine and four for spermine (**Supplemental Fig. S3**). From the HPLC-MS/MS data, it unequivocally supported that the molecular structures of polyamine derivatization products were C_18_H_20_N_2_O_2_ (Put), C_28_H_31_N_3_O_3_(Spd) and C_38_H_42_N_4_O_4_ (Spm) in rice samples (**Fig. 5C**). It concluded that this benzoylation reaction occurred at all N-sites and this quantitative analysis method is correct.

**Figure 5.**
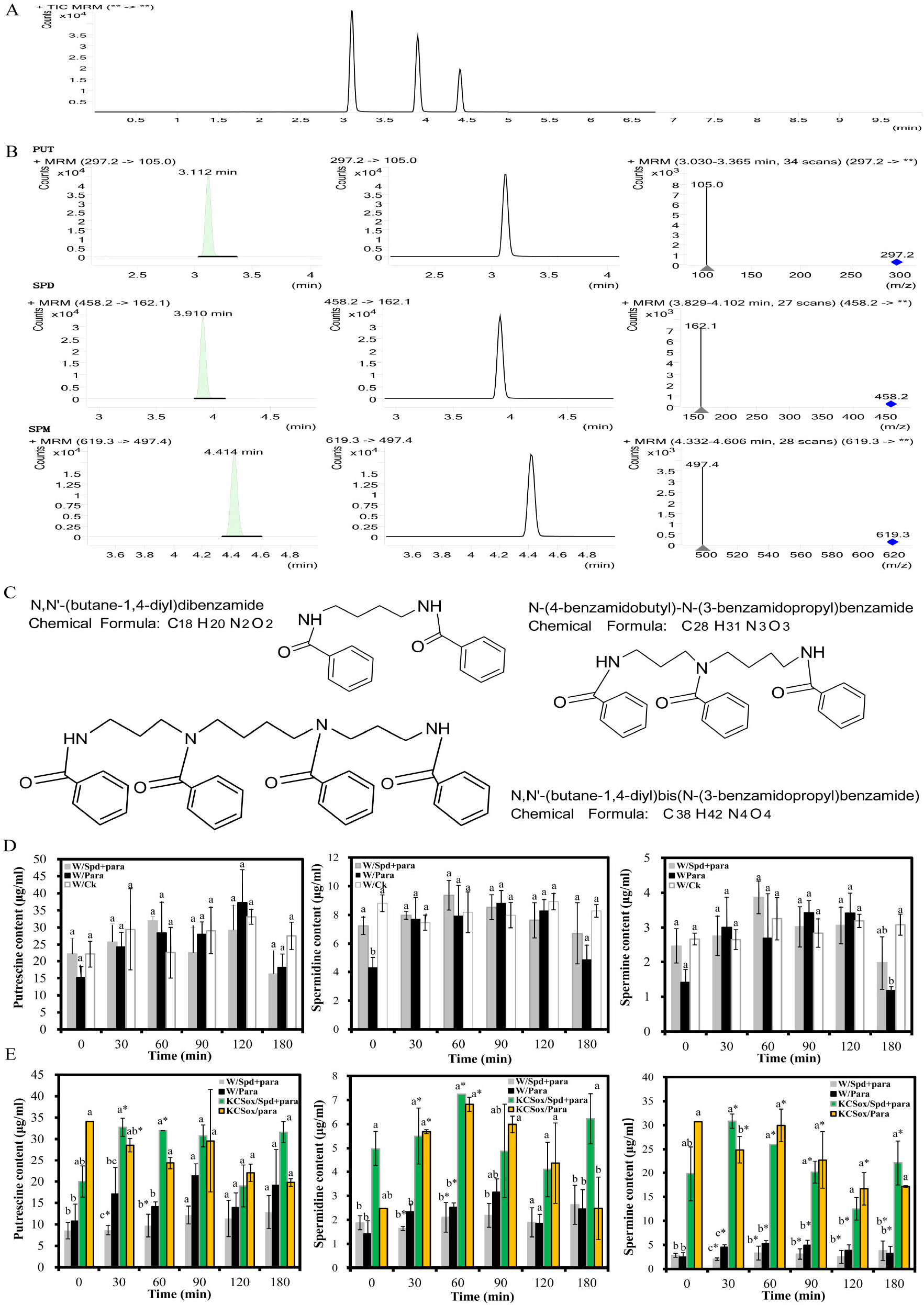
Quantitative analysis of endogenous polyamines in KCSox and WT rice under paraquat and pre-spermidine treatments. **A.** The retention time of standard polyamines derivatizations in the graph of total ions current (TIC) detected by HPLC in order of putrescine, spermidine, and spermine. **B.** The retention time and the exact mass of polyamines derivatizations from rice leaves samples showed in the graphs of multiple reaction monitoring (MRM) detected by HPLC-MS/MS. **C.** Molecular structure of polyamines derivatizations in rice were drawn by ChemBioDraw Ultra with Science of Synthesis Document. **D.** Contents of endogenous putrescine, spermidine and spermine of WT(RHY1179) rice leaves at 6-9 leaf stage in 0, 30, 60, 90, 120 and 180 minutes after sprayed 1.5 mmol/L spermidine 12h before 270mg/L paraquat spraying (Spd+para), 270mg/L paraquat (Para) and ddH_2_O as control (Ck). **E.** Contents of endogenous putrescine, spermidine and spermine of KCSox rice (T2 generation) and WT (Zhonghua11) rice leaves at 6-9 leaf stage in 0, 30, 60, 90, 120 and 180 minutes under the same treatments of Spd+para and Para. Samples of KCSox/Spd+para were 4 replicates at each treatment time, and others were 3 replicates. Different letters indicated different significances among treatments at each time (P < 0.05) and an asterisk indicated significant differences (P < 0.01).

Next, endogenous polyamines contents of KCSox and WT rice leaves at 6-9 leaf stage were quantitatively examined under treatments of pre-spermidine and paraquat (Spd+para), paraquat (Para) and ddH_2_O (Ck). Here it merely observed the contents of spermine in WT rice decreased on 180 min under Spd+para and Para treatments (P<0.5) (**Fig. 5D**), which indicated that all treatments were not critical in contents of polyamines in rice leaves. However, the contents of all three polyamines significantly increased in the KCSox rice than in WT rice under the same treatments (P<0.1) (**Fig. 5E**). This points to the fact that the *EiKCS* gene is related to polyamine biosynthesis. Furthermore, the average contents of polyamines in KCSox rice higher 1.64-fold (Put), 2.02-fold (Spd), and 5.86-fold (Spm) under Para while these contents higher 2.62 times (Put), 2.67 times (Spd), and 7.51 times (Spm) under Para+spd, compared with the WT rice. It supported that the pre-applied exogenous spermidine enhanced the *EiKCS* gene function to promote endogenous polyamines in rice leaves under the paraquat treatment. In addition, the specific contents of all three polyamines in KCSox rice significantly increased at different times. Putrescine and spermidine significantly increased in 30-60 min but spermine increased in 0-180min (P<0.01). Consistent with this, the contents of spermine remarkably increased to a high level up 30μg/ml in KCSox under both Para or Spd+para treatments rather than the level in WT below 5μg/ml. Thus, the prolongation of the overproduction of spermine in KCSox rice was explained by the transformation of endogenous Putrescine and spermidine to spermine in the plant polyamine pathway.

### Overexpression of *EiKCS* gene enhanced antioxidant capacity in rice

To explore the mechanism of paraquat tolerance in the KCSox rice, the variations of physiological indexes were detected about the ROS system, MDA, and Fv/Fw by using the same materials of the same treatments in endogenous polyamines assay. Under the paraquat treatment, the contents of total protein in KCSox rice significantly decreased in 0-180 min, compared to WT rice merely decreasing at 180 min (**Fig. 6A and G**). Moreover, the lipid peroxidation product in KCSox rice leaves induced by paraquat was evaluated in terms of the MDA level. The KCSox rice showed the content of MDA instantly increased 351.62-fold and subsequently decreased in 90 min, although no changes in MDA level was detected in WT rice (**Fig. 6B and H)**. The MDA accumulation may be explained by the role of *EiKCS* itself on the very-long-chain fatty acids (VLCFAs) biosynthesis. Besides, the chlorophyll fluorescence parameters (Fv/Fm) in KCSox rice higher than in the WT rice under paraquat treatment, which indicated *EiKCS* regulated a positive light-harvesting in photosynthesis ((**Fig. 6C and I**). As the biomarker of the antioxidant defenses, the activity per unit protein of SOD, POD, and CAT in KCSox rice showed a great increase throughout the trial period compared with the WT rice (**Fig. 6 D-F and J-L**). In particular, the activities of SOD and POD increased significantly with 12.52-fold and 7.39-fold at 60 min and 44.64-fold and 28.40-fold at 90 min (P<0.01) (**Fig.6 J and K**). These results clearly demonstrated that the overexpression of the *EiKCS* gene in rice could combat the oxidative stress induced by paraquat. Under the spermidine pretreatment, the activities of enzymes and the contents of MDA in KCSox rice had significant increases at 30 min than at 90 min. Again, the pre-applied exogenous spermidine could alleviate the damage of paraquat to both the KCSox rice and WT rice by ahead.

**Figure 6.**
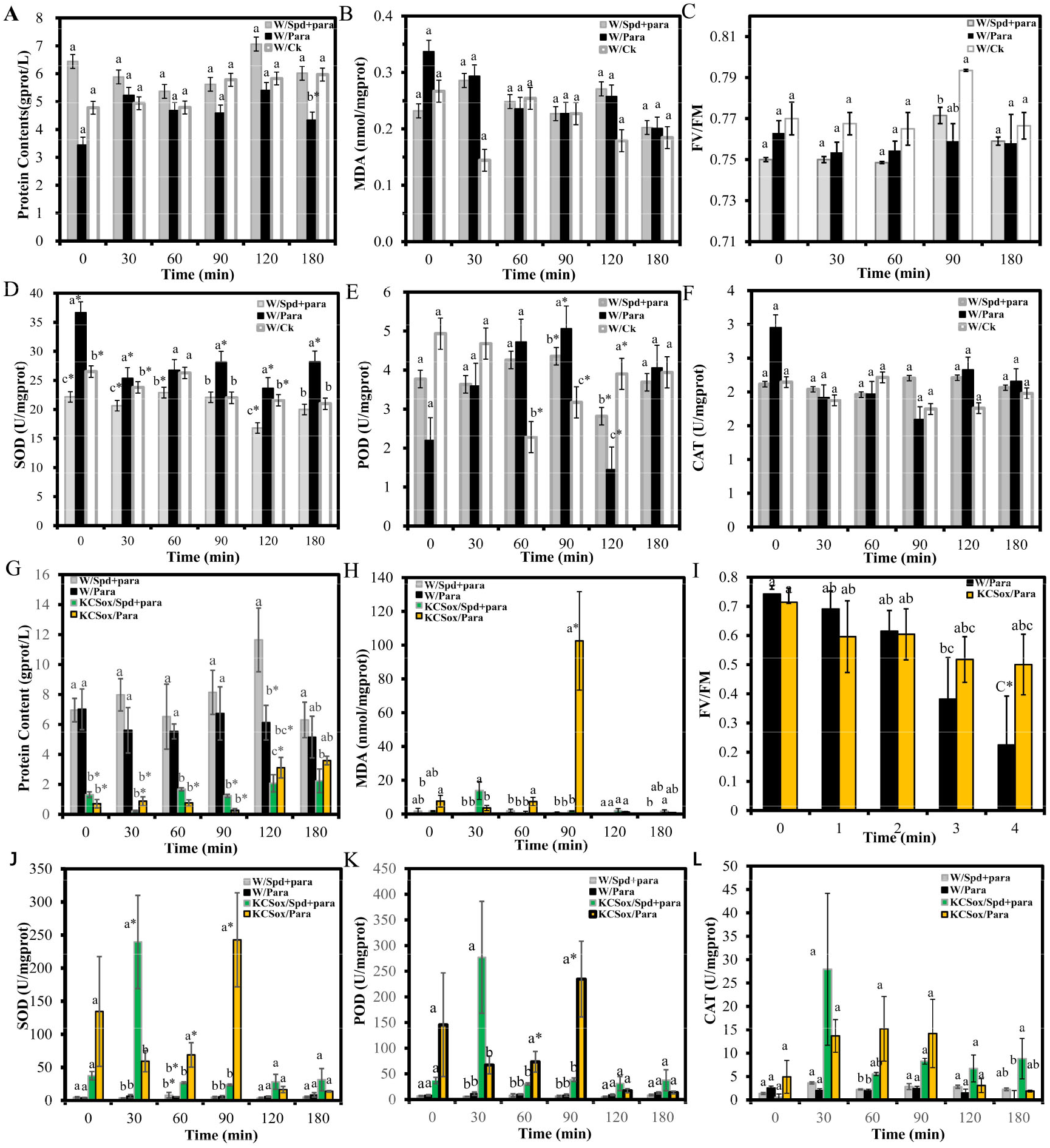
Variations of physiological indexes in KCSox and WT rice under paraquat and pre-spermidine treatments. **A, B and D-F.** the content of total proteins and MDA, and the activities of SOD, POD and CAT in the WT(RHY1179) rice at 6-9 leaf stage in 0, 30, 60, 90, 120 and 180 minutes after sprayied 270mg/L paraquat (Para), 1.5 mmol/L spermidine 12h before 270mg/L paraquat spraying (Spd+para), and ddH_2_O as control (Ck). Samples in WT(RHY1179) rice were 3 replicates at each treatment time. **C and I.** Chlorophyll fluorescence (Fv/Fm) of WT(ZH11) and 3 lines of KCSox(T0) at 4-5 leaf stage with smeared 10mg/L paraquat solution (Para1), 1mmol/L spermidine solution immediately after 10 mg/L paraquat solution (Spd+para1) and ddH_2_O (Ck1) as control for 0, 30, 60, 90, 180 minutes and 0,1,2,3,4 days. Samples in KCSox and WT(ZH11) rice were 3 replicates at each treatment days, and those in WT(ZH11) were 2 replicates at 0, 30, 60, 90, 180 minutes. **G, H and J-L.** The content of total proteins and MDA, and the activities of SOD, POD and CAT in both KCSox (T2 generation) and WT(ZH11) rice at 6-9 leaf stage at 0, 30, 60, 90, 120 and 180 minutes after sprayed 270mg/L paraquat (Para) and 1.5 mmol/L spermidine 12h before 270mg/L paraquat spraying (Spd+para). Samples in KCSox rice were 4 replicates and in WT(ZH11) rice were 3 replicates at each treatment time. All different letters in every graph indicated different significances among treatments at each time (P < 0.05) and an asterisk indicated significant difference (P < 0.01).

### Identification of the EiKCS protein and its cofactors CERs in rice

The mechanism of paraquat tolerance in KCSox rice was further analyzed by proteomics to clarifying these changes in physiological indexes. The proteomic analysis was performed on KCSox and WT rice leaves at 6-9 leaf stage (each 3 samples) under paraquat treatment, in which samples were collected at 48 hours after spraying 270mg/L paraquat. Generally, the whole protein 2605.7μg was extracted from the mixed samples of the 6 rice samples, and then it was analyzed by a label-free qualitative proteomics research (**Supplemental Fig. S4A and Table. S1**). The result showed that 4035 proteins were identified. And the subcellular localization of identified proteins was predicted and classified to mainly gather in the chloroplast (43.82%), cytoplasm (25.53%), and nucleus (14.47%) (**Fig. 7A**). Since paraquat is a PSI inhibitor by utilizing energy through complex enzymes and subcellular fraction of chloroplast, paraquat indeed caused the impact on the function of photosynthesis in energy capture and electron transport-chain maintenance in this study (**Fig. 7B and 7C**). The Lhca1-5 and Lhcab1-6 proteins of the peripheral antennas of PSII and PSI involved in LHCII and LHCI were all identified, which may explain the significant differences of Fv/Fm between KCSox and WT rice leaves under paraquat stress (**Fig. 7B**). And the PsaA-H, PsaK-L, and PsaN-O as 12/16 subunits of the core complex of PSI with critical roles in its function were identified, which may suggest a different excitation energy distribution between PSII and PSI under paraquat stress. Due to ROS burst in chloroplast through pseudo cyclic electron transfer when the free electrons are not sufficient to synthesize ATP or NADPH, the proteins of PetB-C, PetE-H, F-type ATPase and Psb subunits were also identified. These proteins could indicate they protect photosynthetic organs of rice green tissues to related proteins against the paraquat stress-response of ROS from the source (**Fig. 7C**). Consistent with the measured phenotypes of KCSox, it may contribute to promoting the ability of KCSox for enzyme activities in ROS scavenging system.

**Figure 7.**
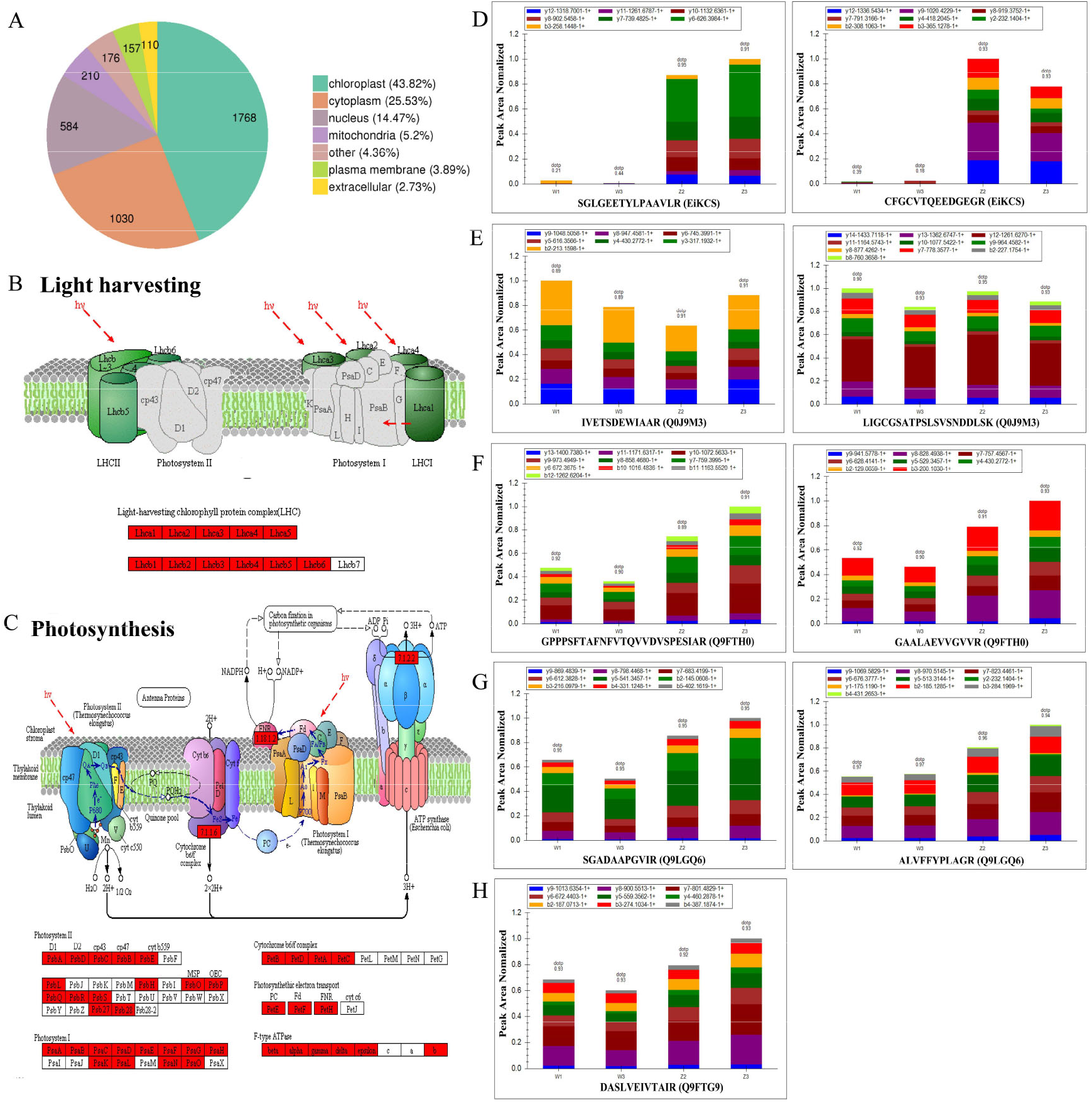
The identified proteins induced by paraquat in qualitative proteomic analysis of mixed samples of rice leaves. **A.** Subcellular localization chart of identified proteins. **B.** Schematic representation of light-harvesting as influenced by paraquat in rice leaves. **C.** Schematic representation of photosynthesis in photosystem I and II as influenced by paraquat in rice leaves. **D.** The distribution of fragment ion peak area on identifications of the EiKCS protein by two synthetic peptide segments. W indicated the WT rice. Z indicated the KCSox rice. **E.** The distribution of fragment ion peak area on identifications the homologous rice proteins (Q0J9M3) of the EiKCS protein by two unique peptide segments. **F-G.** The distribution of fragment ion peak area on identifications the homologous rice proteins (Q9FTH0, Q9LGQ6 and Q9FTG9) of the cofactor CERS protein by their unique peptide segments separately.

However, it cannot be identified that the protein EiKCS in rice corresponding to the overexpression *EiKCS* gene from goosegrass species. Thus, the non-isotopic labeled peptides for EiKCS protein were designed and synthesized according to the protein sequence encoded by *EiKCS* gene **(Table 1)**. Given the outcomes in this study, two of unique peptides SGLGEETYLPAAVLR and CFGCVTQEEDGEGR were confirmed to be used for specific and quantitative detection of EiKCS protein content **(Fig. 7D)**. And all the homologous rice proteins (Q2R3A1 and Q0J9M3) were blasted out from on NCBI website (https://www.ncbi.nlm.nih.gov/) according to the protein sequence of EiKCS protein, the protein Q0J9M3 were found the consistent protein with compared proteomics results by its two effective unique peptides **(Fig. 7E)**. Similarly, the three cofactors CERs of EiKCS protein were identified in this study, which were Q9FTH0, Q9LGQ6 and Q9FTG9 **(Fig. 7F-H)**, according to the homologous CERs sequence (OsCER2, Accession numbers Os04g0611200) provided by the previous reports (Wang et al., 2017).

**Table 1.** Synthesizing and designing of unique peptide segments of EiKCS

Peptide list of protein synthesis of Ei-Ketoacyl-CoA Synthase
|  |  |
| --- | --- |
| 1 | LTGSFTDASLDFQR |
| 2 | SGLGEETYLPAAVLR |
| 3 | CFGCVTQEEDGEGR |
| 4 | VGVSLSR |

### Polyamine pathways were promoted in KCSox rice under paraquat stress

To strengthen clarifications on the function of polyamines in this mechanism of paraquat tolerance in the KCSox rice, quantitative proteomics was performed to detect the expression levels of 15 target proteins associated with the polyamine pathway and other cross-talk pathways. At first, each 50 μg of total protein was separately extracted from 6 rice samples of WT and KCSox (each 3 samples) under paraquat stress as mentioned above, and then they were analyzed by the PRM (Parallel Reaction Monitoring) quantitative proteomics research (**Supplemental Fig. S4B**). After a preliminary evaluation, 2 proteins related to the synthesis and metabolism of polyamine were selected for detection, including the EiKCS protein and three cofactors CERs (**Table 2**). The results showed that 15 proteins of them had evident changes in the levels of expressions under the paraquat treatment **(Fig. 8A)**. The content of EiKCS was higher 51.81 folds in the KCSox rice compared within the WT rice, suggesting that the *EiKCS* gene do overexpress at the protein levels in KCSox under paraquat stress. And high increases in the levels of Q9FTG9, Q9LGQ6, and Q9FTH0 proteins were detected on this KCSox rice, which indicated that the three cofactors were positive correlation with EiKCS in their function of assisting EiKCS catalyzing VLCFAs to wax. Meanwhile, some of the key proteins (Q7X7N2, Q93XI4, and Q9SMB1) regulating polyamines synthesis were also increased the expression levels, which directly promoted the formation of putrescine and its conversion to spermidine and spermine. However, the proteins Q69P84, Q9FRX7, B9F3B6, and O04226 mainly associating with the metabolism of putrescine and spermidine were enhanced while other proteins Q6KAJ2, B9FK36, and Q0J9M3 linking polyamines metabolism to EiKCS via the Tricarboxylic Acid Cycle (TCA) pathway and fatty acid pathway was weakened. Hence, a mechanism for identifying protein-protein interaction with EiKCS were needed to be illustrated by combining 15 proteins with their enriched pathways.

**Table 2.**
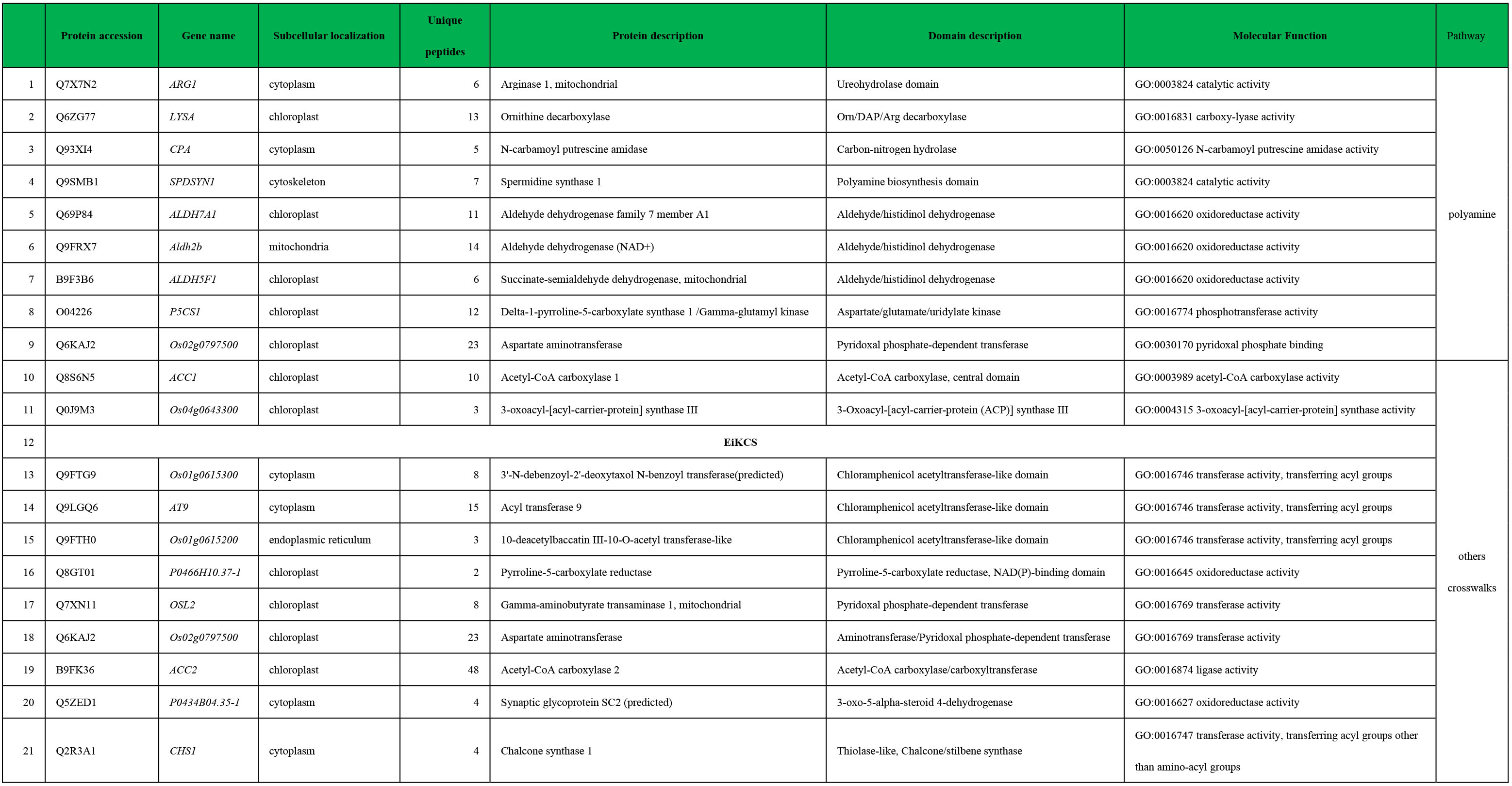
Information of all 21-targeting proteins with functional description

**Figure 8.**
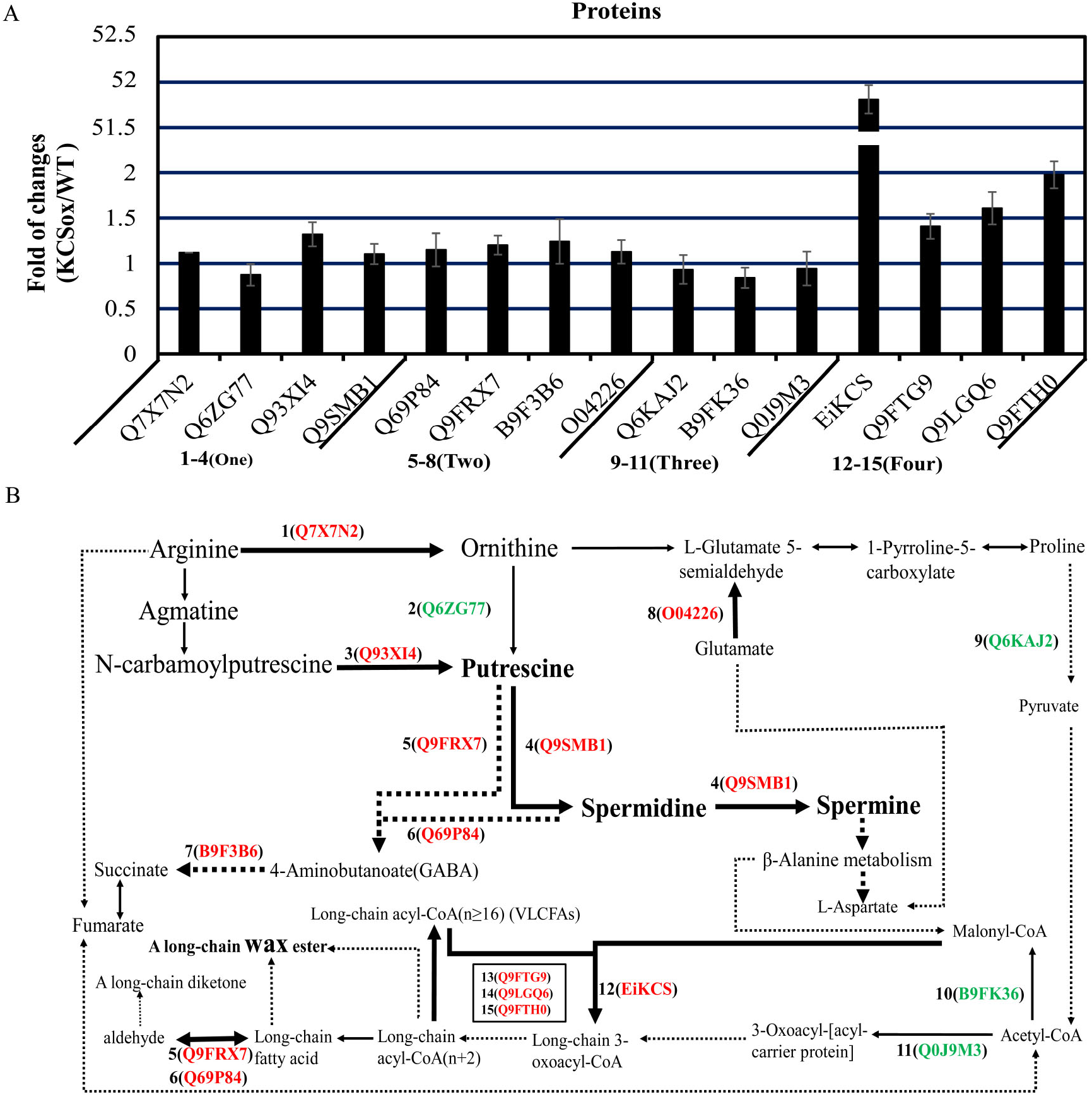
Protein expressions in polyamines pathways and cross-walk with associated pathways. **A.** The changes in the levels of 15 selected target proteins detected by mass spectrometry-based PRM quantitative proteomics in the KCSox and WT rice at 6-9 leaf. The results were presented as a base 2 logarithmic value of the ratio of KCSox/WT (3 replicates). Results of detectable proteins in One group belonged to the synthesis of polyamine pathway, and in Two group belonged to the metabolism of polyamine pathway, and in Three group belonged to TCA pathway and fatty acid pathway, and in Four group belonged to VLCFAs pathway. **B.** The schematic representation of those15 selected target proteins in a path diagram of polyamines pathways and cross-walk with pathways. Red marked proteins indicated up-regulation combined with a bold indicator line. and green marked proteins indicated down-regulation combined with a thinning indicator line. The solid line represents direct regulation and the dotted line represents indirect regulation.

The schematic representation was drawn as a map by combined 15 proteins with their Kyoto encyclopedia of genes and genomes (KEGG) pathway database to illustrate the changes of polyamines related proteins interacting with EiKCS **(Fig. 8B)**. It showed that the overexpression of EiKCS promoted the synthesis of putrescine by the arginine decarboxylase (ADC) pathway rather than the ornithine decarboxylase (ODC) pathway in rice. Especially Q9SMB1 (Spermidine synthase I), it promoted the unidirectional transformation from putrescine to spermidine and then spermidine transferred to spermine, which was consistent with the overproduction of endogenous polyamines contents in the KCSox rice. And the metabolism of putrescine and spermidine separately enhanced by up-regulating Q9FRX7 and Q69P84, which was able to explain the first increase and then decrease of spermidine and the delayed accumulation of spermine. Moreover, the proteins Q9FRX7 and Q69P84 also had a function on fatty acid degradation pathway to directly regulated long-chain fatty acid to aldehyde, which may explain a transient increase of MDA contents in rice under paraquat stress. In sum, overexpression of EiKCS protein via promoting the polyamine pathway and inhibiting the links with the TCA pathway and fatty acid pathway to responding to the paraquat tolerance in the KCSox rice.

## Discussion

The screening process of all offspring of KCSox rice was simplified by those methods in this study with both detection of phenotype and molecular levels. On the one hand, the TPS method reduced the workload of extracting DNA from a large number of leaves samples, and the GMO seeds was soaked in the 25mg/L hygromycin solution for 2 days could eliminate the interference of false-positive in GMO progenies with the hygromycin marker gene. On the other hand, a same vector primer pCUbi1390-F/R could distinguish different genes in MCS of pCUbi1390 vector from positive mono-clones of *E. coli* to T2 generation in GMO seedings, which was more convenient in molecular detection. Moreover, although all KCSox rice were used with the zhonghua11 (ZH11) rice as WT for control to eliminate interferences of original background materials, a commercial rice Ruanhuayou1179 (RHY1179) also were used to explore the function of exogenous spermidine on alleviating paraquat paraquat stress. So those optimal treatment conditions were able to be used in the field production management of other kinds of rice.

On the basis of Naka’s report on quantitative analysis of plant polyamines (Naka et al., 2010), we further showed the molecular formula of the dominant presences of polyamines benzoylation derivatizations in rice. The *EiKCS* gene overexpression significantly increasing endogenous polyamines in rice wouldn’t only be used for enhancing paraquat-tolerance, but also for potential medical and industrial productions, which was supported by a *KCS* gene from *Cardamine graeca* heterologous expressing in *Brassica oilseeds* to engineer high nervonic acid oils for improving human and animal health use (Taylor et al., 2009). And the relationship of EiKCS protein with three cofactors CERs in rice added more information to previous researches on the function of CER for elongation of VLCEAs on other species (Haslam et al., 2012, 2015; Pascal et al., 2013, 2019). Moreover, most proteins of rice were identified in chloroplast under paraquat treatment, which was consistent with the active principle of bipyridine herbicides. After comparison with a similar study, it may be supposed that proteins of photosynthesis and light-harvesting would made a greater impact on the KCSox rice than in a mesosulfuron-methyl resistant *Alopecurus aequalis* (Zhao et al., 2019). Furthermore, the mechanism of paraquat-tolerance in the KCSox rice was also clarified by extending the metabolic pathway of polyamines in this study, which put a new sight to improve the previously reported map of polyamine metabolism and interaction with other metabolic routes (Alcázar et al., 2010).

In this study, the same materials were used to detect the contents of endogenous polyamines and the activities of antioxidant system enzymes, which were mutually connected to the results of proteomics under the same paraquat treatments. Among them, the results of Coomassie brilliant blue method in determining the whole protein concentration in physiological indexes were consistent with the results of BCA protein detection kit, another method in proteomics, which showed average protein content of KCSox (Z2, and Z3) with 2.8350μg/μL was also lower than that of WT (W1, W2, and W3) with 3.7033μg/μL(**Supplemental Table. S2**). Unexpectedly, the line of KCSox (Z1) showed that the *EiKCS* gene was not overexpressed in rice for low contents of EiKCS protein detecting in distribution of fragment ion peak area of the Z1 line (**Supplemental Fig. S5**), although it was indeed a KCSox line after our PCR detection. Therefore, the data of fragment ion peak area distribution and PRM analysis was 2 replicates excluding the replicate of Z1 line and a corresponding W2, and the data with 3 replicates were shown in the supplemental materials (**Supplemental Fig. S6 and Table S3**). The values of two and three replicates didn’t affect the trend of the protein expression in the results that an obvious increase of protein expression in polyamine pathway in the KCSox compared to WT, which was much higher than the fold of change in proteins detected by the same quantitative proteomics (Li et al., 2016).

It is the latest study on polyamines biosynthesis and pathway manipulated in cereals using genetic engineering since it was reported 20 years ago (Capell et al., 1998). Furthermore, this study provided a development of polyamines functions to alleviate paraquat toxicity in GMO crops and paraquat-resistant weed control in crop fields. Thus, it is conclusively that not only can the mechanism in this study contribute to the development of abiotic stress-tolerant GMO crops but also that there is a direct involvement in the solution of multi-resistance in the future herbicide industry. And it can support more studies to investigate the role of EiKCS in goosegrass and subsequent effects in paraquat resistance from both gene overexpressing and silencing.

## Conclusions

The *EiKCS* gene from a resistant-weed goosegrass was copied and transformed into cereal crops using rice as a model and was overexpressed and characterized as an herbicide-tolerance gene, which was demonstrated that *EiKCS* conferred tolerance to paraquat by promoting the synthesis of endogenous polyamines and the expression of proteins in polyamines pathways.

## Materials and methods

### Goosegrass planting and paraquat treatment

A susceptible biotype of goosegrass *(Eleusine indica* L.) was collected from the campus of South China Agricultural University (113°36′, 23°16′N). The resistant (R) biotype of goosegrasswas collected from the Teaching and Research Farm (113°409′, 22°809′N) in Panyu District of Guangzhou. They were also the materials used for a transcriptome profiling and was cultivated by the same methods followed that previous paper (An et al., 2014). In this study, we divided 24 pots with each pot containing 4 plants goosegrass into 3 group. The aboveground parts of every pot as in groups were collected separately. Group 1,2 and 3 with each 8 samples were respectively collected in 30, 60, 90 minutes after spraying paraquat (0.6 kg·ai·ha^−1^). One sample from each group was randomly selected to make a mixed goosegrass sample under paraquat-treatments, and replicated this three times. The methods of spraying paraquat and collecting samples was as same as described in another paper (Luo et al., 2019) (**Supplemental Fig. S7**).

### Full sequence cloning of *EiKCS* gene and expression vector construction

Due to both goosegrass and rice are monocotyledons, *EiKCS* overexpression were constructed in a modified pCUbi1390 vector. The vector Ubiquitin intron and promoter from *Zea mays* in pCUbi1390*::EiKCS* vector is suitable for monocotyledons (Li et al., 2016). We sequenced the pCUbi1390 vector by using a designed primer called Nos-R (5′-ATGTATAATTGCGGGACTCT-3′), and results showed sequence of vector appeared no change. Then three separate RNA samples from leaves of 3 mixed-susceptible goosegrass under paraquat-treatments were extracted, and total RNA was mixed and reverse transcribed to obtain a cDNA by using same kits as described previously (Luo et al., 2019) (**Table. 4-6**). The cDNA serves as a template for amplification of the *EiKCS* gene (1551bp) by primer PqE-F (5′-AATCAGGTCCATCAAGTGTCA-3′)/PqE-R(5′-GGCGGTGTTATCTTCCCT-3 ‘). CDS sequence of this target *EiKCS* gene recombined into the pCUbi1390 vector digested with BamHI and Spel (1563bp). PCR was conducted in a 50μL volume, consisting 2μL cDNA, 2μL of each primer,1μL (25mM each) dNTP, 0.4μL Pfu polymerase and 5μL 10X Pfu buffer (Sangon Biotech, Order NO. B500014). PCR was run in a SimpliAmp™ Thermal Cycler (Thermo Scientific) with the following profile: 95°C 3min, 35 cycles for 94°C 30s, 58°C 30s and 72°C 120s, followed 72°C 6min. And 20μL target KCS fragments in PCR purified products by gel extraction (Sangon Biotech, SanPrep Column PCR Product Purification Kit), 1μL (10μ/μL) each QuickCut™ BamH I /SpeI and 5μl or 10μl Buffer (Takara Code No.1605/1631), 1.5μg pCUbi1390 vector mixed together to form two reaction systems (50μL) which were put into a 37°C constant temperature water bath pot for reaction for 3h, and two target bands was recovered by electrophoresis and gel extraction. Then bands including 6μL (50ng) *KCS* gene from goosegrass and 1μL (100ng) pCUbi1390 vector were mixed with 1μL (5μ/μL) T4 DNAligase, 2μL 10X T4 DNAligase Buffer to total 20 μL in reaction at a constant temperature 22°C for 3h.

### Rice transformation and regeneration

Overexpression pCUbi1390*::EiKCS* vector were transferred into *Escherichia coli (E.colĩ)* DH5a, positive clones were screened in LB plate with Gentamicin (50mg/ml) and Kanamycin(50mg/ml) solutions. Mono-clones are selected added in a same 2ml LB liquid for culture while 1ul bacterial solution of each clone was separately as a template in PCR reactions amplified by designing a primer pCUbi1390-F (5′-TTTAGCCCTGCCTTCATACG-3′)/pCUbi1390-R(5′-TTGCGGGACTCTAATC ATAA-3′) from Ubi-promotor to Nos poly-A (right) including whole sequence of MCS. After the amplified fragments of PCR products were sent for sequencing, monoclonal bacterial solution conforming to the target sequence were used for mass culture. Plasmids in *E. coli* were extracted by the kit by HighPure Maxi Plasmid Kit (TIANGEN Biotech (Beijing) Co.,Ltd., No.DP116).Then this expression cassettes pCUbi1309*::EiKCS* were were introduced into *Agrobacterium tumefaciens (A. tumefaciens)* strain EHA105 by electroporation(1800V/5 ms). Positive clones were screened in LB plate with Hygromycin(50mg/ml) and Rifampicin(20mg/ml). Identification and sequencing of mono-clones in *A. tumefaciens*) also used the primer pCUbi1390-F/R, and their culture is as above. Callus was induced from mature rice embryo of the male-fertile *japonica (O.sativa* ssp. *Japonica* L.) variety Zhonghua11(ZH11), which is a the commonly used varieties for rice transgenic functional verification(Luo et al,. 2013; Wang et al.,2015). *A. tumefaciens-mediated* rice transformation was performed as described (Hiei et al., 1994, 2008).

### KCSox cultivation in fields and screening

To screen positive transformants in each generation rice, cetyltrimethylammonium Ammonium Bromide method (CTAB) was used to extract leaves of the T0 generation rice genome. PCR amplification was used to detect the presence of a hygromycin resistance gene *(hptII)* in genomes of T0 generation rice by amplifying a 289bp sequence with the primer Hyg-F (5′-ACGGTGTCGTCCATCACAGTTTGCC-3′) / Hyg-R (5′-TTCCGGAAGTGCTTGACATTGGGGA-3′). A new TPS method was used to extract DNA from a large number of leaves samples in T1 and T2 generation rice. PCR amplification in T1 and T2 generations rice needed to use both primer pCUbi1390-F/R and Hyg-F/R to detect the presence of pCUbi1390::EiKCS. In order to simplify the detection process, a Fastest DNA Extraction from plant followed by PCR was used to screen positive transformants in T3 generation rice by TIANcombi DNA Lyse&Det PCR Kit (Cat. no. KG203Tiangen biotech(beijing) Co., LtD.). PCR amplification needed to use to detect the presence of pCUbi1390::EiKCS by the primer pCUbi1390-F/R and amplifying a 1468bp sequence with a new primer PqE-1-F(5′-ATTGCTAACTTGCCAGTGTTTCTC-3′)/PqE-1-R(5′-ACGACCAGGA TGCCGATGT-3′).

To use for breeding in fields, we selected some KCSox seedings (T0 generation) in cultivation with ddH_2_O in a greenhouse at 34°C /28°C 12 h (day/night) during 30 days, which is got in 2016 year. After that they (T0) are were planted huanong, campus of South China Agricultural University (113°36′, 23°16′N) from March to November (6 months) in 2017 year (6 months). Seeds of T0 generation rice collected by per plant independently. Then germination and seedling raising of all transgenic rice seedlings (T1) with ddH_2_O in a glasshouse as above. All of rice seedlings (T1) and a new batch of rice seedlings(T0) planted in fields from January to April in 2018 year (4 months) at South breeding and scientific research base of Wuhan University, Lingshui, Hainan (109°45′-110°08’E,18°22’-18°47’N). Seeds of T1 generation rice collected by per plant independently. Germination and seedling raising of some transgenic rice seedlings (T2) with ddH_2_O, and all of rice seedlings (T2) planted in fields from June to November in 2018 year (5 months) at Teaching and research base of South China Agricultural University, Zengcheng, Guangzhou (113°61′-113°81′E, 23°09′-23°13′N). T2 generation rice seeds collected by per plant independently. Then rice seedlings (T3) planted in potted in a greenhouse of campus in South China Agricultural University for experiments in 2019 year (**Supplemental Fig. S8**).

### Grow assay on hygromycin, spermidine and paraquat treatments in the WT rice

“Ruanhuayou1179” (RHY1179) is a kind of commercial hybrid rice (Guangdong Huanongda Seed Industry Co., Ltd), which is suitable for planting in Guangdong province and other southern regions of China with average whole growth period 112-114 days and date of manufacture about these seeds we used is May 21, 2018. Seeds soaked in ddH_2_O for 2 days and germinated on conditions of wet, dark and 35°C. Then seeding were planted in 90 bowls containing regular soil as 45 pots with the same growing conditions of a greenhouse at 34°C /28°C 12 h (day/night). The 6-9 leaf stage of WT (RHY1179) was divided into 30 groups in total with each group contains 3 pots (10 rice seedlings per pot). In order to explore the best screening dose and evaluate their effects on rice growth, 9 groups WT (RHY1179) were firstly sprayed hygromycin solutions with 0, 25, 50, 75, 100, 200, 400, 800 and 1600 mg/L. Secondly, 6 groups were sprayed spermidine solutions with 0, 0.5, 1.0, 1.5, 3.0 and 8.0 mmol/L. Thirdly, 7 groups were sprayed paraquat with solutions 0, 10, 30, 90, 270, 810, 2430mg/L. Precisely, 8 groups were sprayed paraquat with solutions 0, 90, 180, 270, 360, 450, 630 and 810mg/L. This concentrations gradient of hygromycin solutions came from an original liquor of 50 mg·L^−1^ hygromycin solution diluted with ddH_2_O (Brand of Genview, CAS Number 31282-04-9, >80.0%, >1000U/mg, USP, 1g (50mg/ml), Beijing Dingguo Changsheng Biotechnology Co., Ltd).This concentrations gradient of spermidine solutions came from spermidine powder dissolving with ddH_2_O (CAS Number 334-50-9, ≥99.0% (AT), Sigma-Aldrich, SKU-Pack Size 85580-5G). These concentrations gradients of paraquat solution came from a 200 g/L paraquat agent respectively diluted with ddH_2_O (Syngenta, China, as Luo et al., 2019). The spray tower is 3WP-2000 (Nanjing Research Institute for Agricultural Mechanization, Ministry of Agriculture, Nanjing, China). We observed the reaction time after spraying and took photographs as a record and finally measured the above-ground biomass of each pot when group the highest concentration no longer changed after leaf wilting (on 7th days).

Moreover, we confirmed the optimum time of spermidine pretreatment with paraquat. The 6-9 leaf stage of WT (RHY1179) in 18 plastic buckets (r=10cm) of regular soil (each bucket containing 10 plants) were placed in an outside platform outside platform of a particular area in teaching building of South China Agricultural University. They were divided to 6 groups for 6 treatments, which were: (1) spray ddH_2_O as control (2) spray 270mg/L paraquat 24 hours after 1.5 mmol/L spermidine spraying, (3) spray 270mg/L paraquat 12 hours after 1.5 mmol/L spermidine spraying, (4) spray 270mg/L paraquat immediately after 1.5 mmol/L spermidine spraying, (5) spray 270mg/L paraquat 12 hours before 1.5 mmol/L spermidine spraying, (6) spray 270mg/L paraquat 24 hours before 1.5 mmol/L spermidine spraying. The above-ground biomass of each group was weighed separately in the 7th day after its start of spraying (**Table. 7**).

### Germination assay on hygromycin treatments in KCSox and WT

KCSox seeds (60 grains) of T2 generation and WT (RHY1179) seeds (60 grains) were respectively soaked in 9 different concentrations of hygromycin solution as above (0, 25, 50, 75, 100, 200, 400, 800, and 1600 mg/L) for 2 days in a 35°C incubator. Then 15 grains of the KCSox and WT (RHY1179) seeds under each treatment were placed on the filter paper in a culture dish for water culture with ddH_2_O, which was easily distinguished by Toothpicks separating them in the middle and KCSox was an oval rice in the left and wild rice was a long-grain rice on the right. These water cultures of each kind of rice repeated three times, namely 3 culture dishes under each treatment with 45 grains. And the remaining 15 grains of each of the KCSox and wild-type rice seeds were planted in the regular soil for soil culture. All of them were put in the same growing conditions of a greenhouse at 34°C /28°C 12 h (day/night) with only ddH_2_O watering, observed the germination, taken photos, and calculated after 7-10 days. Moreover, another germination had been done as a further validation, which was KCSox rice of T2 generation (100 grains) and WT (ZH11) with 100 grains used to geminate by water culture methods after soaking in 25mg/L hygromycin solution for 2 days. Because KCSox and WT (ZH11) are the same genetic background materials, it can eliminate the influence of materials on germination assay results.

### Grow assay on paraquat treatments in KCSox and WT rice

Under the smear method, paraquat solutions of 10, 100, 665, 1330, 2660, 5320, 10640 and 100000 mg/L were used to explore the half lethal dose of rice leaves. The paraquat concentration of 665mg/L equal to 0.6 kg·ai·ha^−1^ as the recommended rate of fields reported in previous study (Luo et al., 2019). These concentrations were respectively diluted by 200 g·L^−1^ paraquat agent (Syngenta, China, as above) with ddH_2_O, and them used by cotton to applied on the leaves of the corresponding rice seedlings. Leaves were of WT (ZH11) at 4-5 leaf stage, which were planted in the plastic bucket (r=8cm) of soil adding 13g compound fertilizer (each bucket containing 5 plants) and were photographed four days later after smearing paraquat at a greenhouse 34°C /28°C 12 h (day/night). Percent inhibition rate of leaves under each paraquat concentration was calculated by the number of dead leaves dividing by randomly smear number of total leaves (**Supplemental Fig. S9 and Table. 8**).

Under the sprayed method, KCSox (T2 generation) and WT (ZH11) at 6-9 leaf stage were sprayed 270mg/L paraquat solutions, which is used to calculate the survival rate paraquat damages on 7th day. At first, T2 generation seeds of three KCSox lines with random selection (each 50 grain) geminated by water culture methods after soaking in 25mg/L hygromycin solution for 2 days. And WT (ZH11) as the same genetic background materials as a control after soaking in ddH_2_O for 2 days. Secondly, they were kept wet on culture-dishes 7 days with ddH_2_O and the germinated seeds from them were selected out and put in the bowl with regular soil (each bowl contain 1 grain) at an outside platform of a particular area in the teaching building of South China Agricultural University for experiments. Thirdly, the penultimate leaf of each rice seedlings was detected by PCR reaction to make sure the presence of the target *EiKCS* gene. Finally, twelve KCSox rice plants (12 bowls) and twelve WT (ZH11) rice plants (12 bowls) with the same growth trend were selected. And then 9 bowls from 12 bowls in each of the KCSox and WT (ZH11) rice were sprayed with the paraquat solutions and those rice in other bowls were sprayed ddH_2_O as the control. And then these seedlings in 18 bowls from 24 bowls of the KCSox and WT (ZH11) rice were sprayed with the paraquat solutions. The other seedlings in the left 6 bowls were sprayed ddH_2_O as the control. After spraying, they were divided into three groups with 4 bowls as a group and were photographed at 0, 12, 24, 36, 48, 60, 160 hours (**Supplemental Fig. S10**). it is until the 7th day when the above-ground biomass of each bowl was separately measured.

### KCSox on paraquat and pre-spermidine with paraquat treatments for detection of polyamines (Put, Spd and Spm), antioxidant enzymes (SOD, POD and CAT) and harmful substances (MDA)

Under the sprayed method, Seeds of WT(RHY1179) wild rice after germination with ddH_2_O were planted in 18 bowls of regular soil (each bowls containing 30 grains). WT (RHY1179) at 6-9 leaf stage of were sprayed 270mg/L paraquat, 1.5 mmol/L spermidine 12h before 270mg/L paraquat spraying, and ddH_2_O as control for 0, 30, 60, 90, and 180 minutes. In order to eliminate the any additional interference, three lines t of KCSox seeds (T2 generation) germinated with ddH_2_O, which was as the same lines in grow assay of paraquat. They were planted in 9 plastic buckets (r=10cm) of regular soil (each bucket containing 30 grains). At 6-9 stage, only half of them are transplanted to bowls with one plant in a bowl. 7 days after transplanting is used to naturally recover the growth of plants. During this period, transgenic detection of *EiKCS* gene was carried out for each plant. Then they are selected into 2 groups. One group is 42 bowls KCSox plants, and the other group is 36 bowls false-positive transgenic rice plants as wild rice for control materials (WT). At 6-9 leaf stage, they are sprayed 270mg/L paraquat and 1.5 mmol/L spermidine 12h before 270mg/L paraquat spraying for 0, 30, 60, 90, and 180 minutes. The aboveground parts of each plant were taken and collected as one sample, which then immediately frozen inliquid nitrogen and stored at −80°C for detection of polyamines, antioxidant enzymes (SOD, POD and CAT) and harmful substances (MDA).

### Polyamines Extraction, Benzoylation, and HPLC-MS/MS

Free Polyamines extraction from plants, benzoylation of the extracted polyamines were performed as described in previous reports with modifications (Naka et al., 2010; Takahashi et al., 2018). In order to determine more accurately, we improve several fixed values. 0.3g plant samples were pulverized into powder with a mortar and pestle in liquid nitrogen environment. Then take 0.3g powder to 10ml centrifuge tube, add 1.5ml of 5% (v/v) cold perchloric acid (PCA), and keep it on ice for 1 h. Centrifugation of 10000×g for 30 min at 4°C. Collect 1.5 ml the supernatants and add 1ml NaOH (2 mol/L) to plant extract, rotate, then add 10 ml benzoyl chloride, mix and culture at room temperature for 20 minutes. It is important to ensure that benzoyl chloride is C_7_H_5_ClO (140.57g/mol, CAS Number 98-88-4, ≥98.0%, Shanghai Lingfeng Chemical Reagent Co., Ltd) because it determines the molecular formula of polyamines derivatizations in the detection. Add 2 ml of saturated sodium chloride. Then 2 ml of ether was added and forced to mix, and 3000×g was centrifuged at 4 *°C* for 10 minutes. Take 1ml of the supernatant and filter it with a filter syringe (aperture 0.2mm). Add the filtrate into a 2ml sample bottle, evaporate the organic solvent phase with hot air, and resuspend the residue in 1ml of methanol.

The benzylated polyamines were analyzed by liquid chromatography-mass spectrometry (HPLC-MS, Uplc1290-6470A QQQ, eclipse plus, 2.1X50mm, 1.8um, Agilent). The mobile phase 0.2% formic acid aqueous solution and acetonitrile were determined. External standard method was used for quantitative analysis of derivatization products of three kinds of polyamine benzoyl chloride. Standards of putrescine, spermidine, and spermine (CAS Number 110-60-1, 124-20-9, 334-50-9, ≥99.0% (AT), Sigma-Aldrich, SKU-Pack Size 51799-100MG, 49761-100MG and 55513-100MG) were used to prepare 1.5ml (1mmol) spermidine, spermine and putrescine solution respectively, then prepare as 2ml of ether phase. Take 0.5ml of each ether phase of polyamines, add in 2ml of sample bottle respectively, and add 1ml of methanol after the ether solution volatilizes. Considering the linearity and the optimum concentration of the sample, three polyamines derivatization putrescine (C_18_H_20_N_2_O_2_, 297>105), spermidine (C_28_H_31_N_3_O_3_, 458>162), spermine (C_38_H_42_N_4_O_4_, 619>497) were quantified by standard curve. The limit of polyamines detection was 5ng/ml and the limit of quantification was 1ng/ml.

### The total protein and MDA contents, and SOD, POD and CAT activities assays

The level of intracellular ROS scavenging capacity and the extent of lipid peroxidation in rice were analyzed in the same materials of polyamines detections. 0.1g plant samples were pulverized into powder with a mortar and pestle in liquid nitrogen environment for detection by each indicator. All indicators were measured with the corresponding detection kit according to the manufacturer’s instructions (Nanjing Jiancheng Bioengineering Institute, Nanjing, China). The total protein content detected by means of Coomassie brilliant blue protein determination kit (A045), which used in calculation of SOD (A001-1), POD (A084-3) and CAT (A007-1) activities and MDA (A003-1) content, so that their determinations were conducted and shown by U·mgprot^−1^(Sun et al., 2013; Shu et al., 2019).

### Maximum quantum yield of photosystem II (Fv/Fm)

To assess the rice photosynthetic performance after exposure to paraquat, we detected the maximum quantum yield of photosystemII (Fv/Fm) according to the previous measurement method (Brunharo et al., 2017). Under the smearing method, leaves of WT (ZH11) at 4-5 leaf stage at a greenhouse 34°C/28°C 12h (day/night) were treated with 10mg/L paraquat solution (para), 1mmol/L spermidine solution immediately after 10 mg/L paraquat solution (spd+para) and ddH_2_O (ck) as control. The flag leaves of rice were randomly selected for Fv/Fm determination with chlorophyll fluorometer (Mini PAM, Waltz, Effeltrich, Germany) at 0, 30, 60, 90 and 180 minutes after smearing (2 blades for each treatment). The treated time 0 minute meant that it was started after each treatment for an hour and a half because they were put in the sunshine for 1 hour for the paraquat reaction and moved into the dark room for 30 minutes for dark measurement environment.

Similarly, we randomly selected T0 generation 3 lines of KCSox and T0 generation 3 lines of false-positive KCSox as control materials (WT) at 4-5 leaf stage at a greenhouse 34°C/28°C 12h (day/night) were treated with 10mg/L paraquat solution (para) under the smearing method. Their flag leaves were detected Fv/Fm at 0, 1, 2, 3 and 4 days after smearing. The treated time 0 day also meant that it was started after the treatment for an hour and a half.

### Transgenic *EiKCS* rice for proteomics analysis

Leaves of three plants of T3 generation KCSox, and leaves of three plants of T2 generation KCSox (Z1, Z2, and Z3), and three plants of WT (ZH11) rice (W1, W2, and W3) at 6-9 leaf stage (from three individual plants each) at a greenhouse 34°C/28°C 12h (day/night) were sprayed with 270mg/L paraquat solution for 2 days. Firstly, leaves from all KCSox were mixed as one sample for label-free quantitative proteomics analysis. Secondly, leaves from Z1, Z2, Z3, W1, W2, and W3 were separately saved as 6 samples for PRM (Parallel Reaction Monitoring) analysis. The leaves were placed in a centrifugal tube and frozen in liquid nitrogen at −80 °C.

### Protein extraction

The samples stored at −80 °C were ground by liquid nitrogen into cell powder and then transferred to a 5mL centrifuge tube. After that, four volumes of lysis buffer (8 M urea, 1% Triton-100, 10 mM dithiothreitol, and 1% Protease Inhibitor Cocktail) was added to the cell powder, followed by sonication three times on ice using a high-intensity ultrasonic processor (Scientz-IID, Ningbo Scientz Biotechnology Co., Ltd. China). The remaining debris was removed by centrifugation at 20,000 g at 4 °C for 10 min. Finally, the protein was precipitated with cold 20% trifluoroacetic acid for 2 h at −20 °C. After centrifugation at 12,000 g 4 °C for 10 min, the supernatant was discarded. The remaining precipitate was washed with cold acetone for three times (All reagents from Sigma-Aldrich and Calbiochem). The protein was redissolved in 8M urea and the protein concentration was determined with BCA Protein Assay Kit (P0009, Shanghai Beyotime Biotechnology CO., LTD, China) according to the manufacturer’s instructions.

### Trypsin digest and peptides synthesis of proteins, especially EiKCS

The protein solution samples were hydrolyzed into peptide segments by trypsin (V9012, Promega (Beijing) Biotech Co., Ltd, China) for label-free quantitative proteomics and PRM analysis. Importantly, the Ketoacyl-CoA synthase protein was encoded by the *EiKCS* gene, four unique peptides of which were selected according to its sequence by Skyline software (v.3.6) for peptide synthesis. After the synthesis, the reductive alkylation experiment was carried out for PRM analysis.

### Label-free quantitative proteomics analysis

The process of all proteins was performed based on the methods (Zhao et al., 2020) with a slight modification of parameters. HPLC Fractionation was used that the gradient of the peptide was 8% - 32% acetonitrile, pH 9, over 60 minutes into separate 60 components, then the peptide was combined into 18 components and freeze-dried in a vacuum. After the peptide was separated by EASY-nLC 1000 UPLC system (Thermo Scientific, Carlsbad, U.S.A) at a constant flow rate of 450 nL/min. The peptides were subjected to NSI source followed by tandem mass spectrometry (MS/MS) in Q ExactiveTM Plus (Thermo Fisher Scientific Inc., U.S.A) coupled online to the UPLC. Automatic gain control (AGC) was set at 5E4. Fixed first mass was set as 100 m/z. Maxquant (v1.5.2.8) was used to retrieve the secondary mass spectrometry data. Retrieval parameter setting: rice Swissprot database (48915 sequences). The false discovery rate (FDR) was adjusted to <1% and minimum score for peptides was set >40. It was also used label-free quantification to determine the normalized protein intensity.

### Subcellular Localization

There, we used WoLF PSORT a subcellular localization predication soft to predict subcellular localization. WoLF PSORT is an updated version of PSORT/PSORT II for the prediction of eukaryotic sequences (https://omictools.com/wolf-psort-tool).

### PRM analysis

To validate the reliability of label-free quantitative proteomics results, a PRM assay was performed using the same six protein samples. After the assessment of primary results, we selected 15 proteins from the polyamine pathway and their associated pathways. These proteins detected by their unique peptides in the PRM assay. Each rice protein has at least 3 unique peptides and the Ketoacyl-CoA synthase protein encoded by the *EiKCS* gene used 4 synthesized peptides. The full process of PRM assay were according to previous details (Zhao et al., 2019) with a little parameter’s modifications. Briefly, After the peptide was separated by EASY-nLC 1000 UPLC system at a constant flow rate of 350 nL/min. The peptides were subjected to NSI source followed by tandem mass spectrometry (MS/MS) in Q ExactiveTM Plus coupled online to the UPLC. The electrospray voltage applied was 2.0 kV. The m/z scan range was 350 to 1080 m/z for full scan, and intact peptides were detected in the Orbitrap at a resolution of 70,000. Peptides were then selected for MS/MS using NCE setting as 27 and the fragments were detected in the Orbitrap at a resolution of 17,500. A data-independent procedure that alternated between one MS scan followed by 20 MS/MS scans. Automatic gain control (AGC) was set at 3E6 for full MS and 1E5 for MS/MS. The maximum IT was set at 50ms for full MS and auto for MS/MS at 120ms. The isolation window for MS/MS was set at 1.6 m/z. Peptide settings: enzyme was set as Trypsin [KR/P], Max missed cleavage set as 0. The peptide length was set as 7-25. The product ions were set as from ion 3 to last ion, the ion match tolerance was set as 0.02 Da.

### Pathway enrichment by KEGG annotation

Kyoto Encyclopedia of Genes and Genomes (KEGG) database was used to annotate protein pathway. Firstly, using KEGG online service tools KAAS to annotated protein’s KEGG database description. Then mapping the annotation result on the KEGG pathway database using KEGG online service tools KEGG mapper. KEGG database was used to identify enriched pathways by a two-tailed Fisher’s exact test to test the enrichment of the identified protein against all proteins of the species database. The pathway with a corrected p-value of <0.05 was considered significant. These pathways were classified into hierarchical categories according to the KEGG website (http://www.kegg.jp/).

### Data analysis

Data were analyzed by analysis of variance (ANOVA) at Duncan’s test a 95% (P = 0.05) and 99% (P = 0.01) confidence level using Excel 2016, and SPSS 17.0 (SPSS Inc., Chicago, IL, USA), and the Processing System (DPS 7.05, Zhejiang, China). For proteins in each category, InterPro (a resource that provides functional analysis of protein sequences by classifying them into families and predicting the presence of domains and important sites) database was researched and a two-tailed Fisher’s exact test was employed to test the enrichment of the identified protein against all proteins of the species database. Protein domains with the P value < 0.05 were considered significant.

## Supplemental materials

**Supplementary Fig. S1** Proteins from transcriptome splicing sequence of the KCS gene in susceptible and resistant biotype goosegrass without paraquat treatment. Results of next-generation sequencing from samples (C_S_comp30798_c0_seq2) susceptible goosegrass seedlings without paraquat (S0); (C_S_comp30798_c0_seq2) resistant goosegrass seedlings without paraquat (R0).

**Supplementary Fig. S2** DNA extraction steps of TPS method.

**Supplementary Fig. S3** Seven possible molecular structures for polyamines derivatizations. Two for putrescine (Put), three for spermidine (Spd) and four for spermine (Spm).

**Supplementary Fig. S4** Quality control of protein extraction. The 6-9 leaf stage of ZH11 wild rice (W) and T3 generation transgenic EiKCS rice (Z). Each 3 biological replicates. **A**. The gel of SDS-PAGE of the mixed rice sample. **B**. The gel of SDS-PAGE of the 6 rice samples.

**Supplementary Fig. S5** Distribution of fragment ion peak area on identifications of the EiKCS protein by two synthetic peptide segments. Each 3 biological replicates. W indicated the WT rice. Z indicated the KCSox rice.

**Supplementary Fig. S6** Changes in the levels of 15 selected target proteins detected by mass spectrometry-based PRM quantitative proteomics in the KCSox and WT rice at 6-9 leaf. The results were presented as a base 2 logarithmic value of the ratio of KCSox/WT (3 Biological replicates and 3 technical replicates).

**Supplementary Fig. S7** Materials for Acquisition RNA and cDNA of EiKCS Gene. Goosegrass *(Eleusine indica* L.) in 24 pots before paraquat treatments.

**Supplementary Fig. S8** *Agrobacterium* mediated transformation of rice and rice cultivation in fields. The information of time and places about transgenic rice is below every picture. T0 generation rice cultivated two kinds of transgenic rice, which is transgenic *EiKCS* rice and transgenic *EiPqTS3* rice (as control). They are separately transformed by pCUbi1390::EiKCS and pCUbi1390::PqTS3 vectors. Then T1, T2 and T3 generation of transgenic rice cultivated only transgenic *EiKCS* rice.

**Supplementary Fig. S9** Photographs of Rice leaves smearing paraquat solutions with different concentrations. Smearing 4-5 leaf stage of wild rice (ZH11) with different concentrations of paraquat solution (smear method). The paraquat concentration of 665mg/L equal to 0.6 kg·ai·ha-1 as the recommended rate. Phenotypes after paraquat treatments 4 days.

**Supplementary Fig. S10** Phenotypes of paraquat damages of KCSox and WT (zhonghua11) rice at 6-9 leaf stage after spraying 270mg/L paraquat solutions, which were photographed at 0, 12, 24, 36, 48, 60, 160 hours. Twelve KCSox rice plants (12 bowls) and twelve WT (ZH11) rice plants (12 bowls) are selected with the same growth trend.

**Table. S1** Protein concentration for label free qualitative proteomics research.

**Table. S2** Proteins concentrations for the PRM (Parallel Reaction Monitoring) quantitative proteomics research.

**Table. S3** The expression levels of 15 selected target proteins detected by PRM quantitative proteomics in the KCSox rice (Z) and WT rice (W).

**Table. S4** Randomly select one of S1, S2 and S3 as a mixing sample for grinding, which replicated 3 times.

**Table. S5** Detection concentrations for RNA extracted of from mixed sample of goosegrass sprayed with paraquat.

**Table. S6** Total RNA was mixed and reverse transcribed to obtain a cDNA of goosegrass under paraquat treatments.

**Table. S7** Determination of optimum time of spermidine pretreatments with paraquat. The 6 9 leaf stage of wild rice (RHY1179) in 18 plastic buckets (r=8cm) of regular soil (each bucket containing 10 plants). (1) spray ddH_2_O as control, (2) spray 270mg/L paraquat 24 hours after 1.5 mmol/L spermidine spraying, (3) spray 270mg/L paraquat 12 hours after 1.5 mmol/L spermidine spraying, (4) spray 270mg/L paraquat immediately after 1.5 mmol/L spermidine spraying, (5) spray 270mg/L paraquat 12 hours before 1.5 mmol/L spermidine spraying, (6) spray 270mg/L paraquat 24 hours before 1.5 mmol/L spermidine spraying.

**Table. S8** Percent inhibition rate of leaves under each paraquat concentration by smearing 4-5 leaf stage of wild rice (ZH11) with different concentrations of paraquat solution (smear method).

## Acknowledgment

This work was supported by the Key Realm R&D Program of GuangDong Provice (2019B020221002); the National Natural Science Foundation of China (No. 31871980) and the Science and technology program of Guangdong Province (2019B121201003-X). We thank for Jan Yan (College of Natural Resources and Environment, South China Agricultural University) for his technical help with molecular formula analysis of polyamine benzoylation derivatives, Shaokui Wang (College of agriculture, South China Agricultural University) for her technical help with TPS method for testing, and Yangsheng Li (College of Life Sciencest, Wuhan University) for his providing help of Lingshui of rice teaching and breeding base, the assistance during experiments preparation of Mengnuan Ran, Yunfei Fu and Chengxi Deng.

